# Electrical propagation of vasodilatory signals in capillary networks

**DOI:** 10.1101/840280

**Authors:** Pilhwa Lee

## Abstract

A computational model is developed to study electrical propagation of vasodilatory signals and arteriolar regulation of blood flow depending on the oxygen tension and agonist distribution in capillary network. The involving key parameters of endothelial cell-to-cell electrical conductivity and plasma membrane area per unit volume were calibrated with the experimental data on an isolated endothelial tube of mouse skeletal feeding arteries. The oxygen saturation parameters in terms of ATP release from erythrocytes are estimated from the data of a left anterior descending coronary blood perfusion of dog. In regard to the acetylcholine induced upstream conduction, our model shows that spatially uniform superfusion of acetylcholine attenuates the electrical signal propagation, and blocking calcium activated potassium channels suppresses that attenuation. On the other hand, local infusion of acetylcholine induces enhanced electrical propagation that corresponds to physiological relevance. In the integration of the endothelial tube and the lumped arterioles vessel model, we show that endothelial purinergic oxygen sensing of ATP released from erythrocytes and local infusion of acetylcholine are all individually effective to induce vasodilatory signals to regulate blood flow in arterioles. We have recapitulated the upstream vasomotion in arterioles from the elevated oxygen tension in the downstream capillary domain. This study is a foundation for characterizing effective pharmaceutical strategies for ascending vasodilation and oxygenation.

## 1 Introduction

We investigate the influence of vasodilatory signals in the electrical propagation in the endothelium of microvasculature and the dilation in the upstream arterioles. Blood flow regulation in the microcirculation is crucial for keeping relevant pressure tone and tissue oxygenation. In arterioles and capillary microvasculature, the downstream signaling of oxygen demand or hypertension induces upstream conducted vasodilation (Bagher and Segal 2011). The upstream propagation of electrical signals in the microvasculature is activated in the vasoregulation in exercise at healthy state (Duncker and Bache 2008), and does malfunction in diabetes (Fitzgerald et al. 2005, Park et al. 2008, Bender et al. 2009), obesity (Haddock et al. 2011), and hypertension (Brahler et al. 2009) at disease states. Microvascular vasodilation is regulated by agonists such as cholinergic acetylcholine (ACh) (Doyle and Duling 1997), ATP (Ellsworth 2000), and also via sympathetic control of *β*-adrenergic signaling (Feigl 1998). Dilation in coronary arterioles is also flow dependent (Kuo et al. 1990). We target the response to agonists of ACh and ATP through a couple of steps (Chen et al. 1992, Communi et al. 2000, Raqeeb et al. 2011) or directly to pharmaceutical drug of SK_Ca_ and IK_Ca_ (small and intermediate conductance Ca^2+^ activated K^+^ channel) opener, NS309 (6,7-dichloro-1H-indole-2,3-dione 3-oxime).

When the mechanisms of the hyperpolarization and vasodilation is briefly summarized, in the initiation, the vasodilatory agonists increase intracellular calcium via G-protein coupled receptor and IP_3_ induced calcium release from endoplasmic reticulum. The elevated calcium hyperpolarizes the endothelial cell (EC) membrane via SK_Ca_ and IK_Ca_ (Stankevicius et al. 2011). The hyperpolarization in EC is transmitted to the smooth muscle cell (SMC) layers via myoendothelial junction, and closes L-type calcium channels, lowering the influx of calcium ions. The elevated calcium in EC also produces nitric oxide (NO), and the increased NO is transported to SMC layers to decrease calcium through cGMP pathway. The conduction of the hyperpolarization and vasodilation is primarily through endothelial GJCs (gap junction channels) (Goto et al. 2002, Diep et al. 2005, Schmidt et al. 2008).

We study the following questions to characterize the vasodilatory signal propagation in model capillary networks: (1.) How far can a localized hyperpolarized signal propagate? (2.) What are the conditions under which a physiologically meaningful depolarization (tens of millivolts) occur (3.) How influential are endogenous agonists and pharmaceutical drugs in the upstream vasodilation? Using our endothelial tube model with an endothelial cell model modified from Silva et al. (2009), we realized the electrical signal propagation speed about 2.5 mm/s and electrical length constant (*λ*) about 1.1 mm that are consistent with the experimental data (Segal and Duling 1986). In model simulation, *λ* decreases with more ACh/NS309 when the dilators are uniformly superfused. In contrast, local application of the agonist makes the electrical signals to propagate further, and indeed this captures some physiological conditions for different effects of ACh to the electrical propagation (Emerson et al. 2002, Behringer et al. 2012).

The integrated model of endothelial tube and lumped arterioles (i.e. endothelial cells are coupled to smooth muscles via myoendothelial junctions) shows physiological vasodilatory response from the downstream low oxygen tension observed in canine coronary blood flow (Farias et al. 2005) or cholinergic signals in skeletal muscle of hamster (Bagher and Segal 2011). Here, the model oxygen sensing is assumed to be from purinergic binding of ATP from erythrocytes (Ellsworth 2000) and to effect on IP_3_ generation rate in endothelium (Communi et al. 2000, Raqeeb et al. 2011). The proposed endothelial tube model in capillary network is expected to provide a foundation for further study in the blood flow regulation from vasodilatory signals from pathphysiological states such as ischemic hypoxia and hypertension as well as metabolic syndrome, and the molecular and cellular studies for potential therapeutic and preclinical drug screening.

## 2 Methods

### 2.1 Mono-domain capillary network

In the mono-domain formulation, the electrical current in the EC layer is represented by the electrical conductivity *σ*_EC_ as follows:

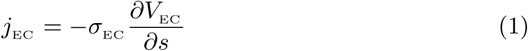

In the continuity equation on the electrical endothelial currents, the current from membrane capacitance per unit area *C*_m_, membrane ionic currents 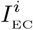, and external current injection *I*_INJECTION_ induce the gap junctional transport in the following:

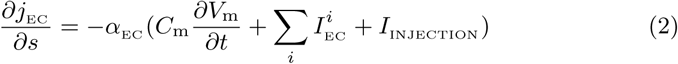

where *V*_m_ is exactly the same as *V*_EC_ with extracelllar matrix grounded as zero. The transmitted electrical currents from the extracellular matrix to the EC layer are explicitly represented with 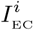 for *i*^th^ ionic species (Na^+^, K^+^, Cl^−^, Ca^2+^). The scaling factor *α*_EC_ is the endothelial plasma membrane area per unit volume. The cellular solute/ionic dynamics basically follows the mathematical formulations in Silva et al. (2009) for ionic currents and transporter/exchangers:

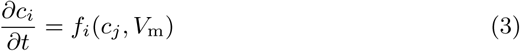

In the consideration of direct agonist NS309 of calcium activated potassium channels (SK_Ca_/IK_Ca_), we formulated the open probability with the competitive binding of [Ca^2+^]_in_ and [NS309] based on the data from Behringer and Segal (2012):

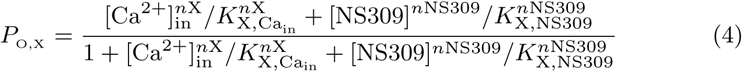

where X indicates SK_Ca_ or IK_Ca_, and *n*X and *K*_X,Ca_ denote the Hill coefficient and the half saturation coefficient for calcium. For indirect agonist of ACh, we changed the IP_3_ generation rate proportional to [ACh], with the IP_3_ generation rate as 6.6 × 10^−8^ mM/ms for [ACh]=3*µ*M. Most parameters are the same as Silva et al. (2007), and the others are enlisted in Table 1. In the capillary network, the boundary condition for downstream terminal nodes is *j*_EC_ = 0 so that the terminal nodes are sealed electrically. On the other hand, boundary conditions for bifurcation nodes are prescribed as Σ_*i*_ *j*_EC_, *i* = 0 satisfying the conservation of currents at bifurcation nodes.

**Table 1.** Endothelial parameters.

| Symbol | Definition | Value |
| --- | --- | --- |
| $\sigma_{EC}$ | EC electrical conductivity | $1.3 \times 10^{-15}$ S cm |
| $\alpha_{EC}$ | EC plasma membrane area per unit volume | 0.32 / cm |
| $C_m$ | Membrane capacitance per unit area | 0.02 pF |
| $g_{SKCa}$ | Maximum $SK_{Ca}$ conductance | 0.1534 nS |
| $g_{IKCa}$ | Maximum $IK_{Ca}$ conductance | 0.4254 nS |
| $K_{NS309}$ | Half saturation coefficient for NS309 | 1.0 μM |
| $n_{NS309}$ | Hill coefficient for NS309 | 1.3 |

In the upstream intact to terminal arterioles, we couple the endothelium with the lumped arterioles in the following:

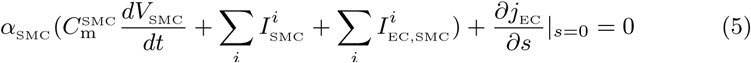

where the parameters for the smooth muscle electrophysiology 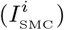 and myoendothelial junctions 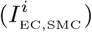 are mostly from Kapela et al. (2008, 2009).

### 2.2 Blood flow perfusion

The blood flow perfusion is determined by the diameter of the lumped pre-capillary arterioles with the assumption that the pressure gradient is constant. We apply the law of Laplace at the equilibrium of vessel mechanics for the determination of arterioles diameter in terms of SMC calcium concentration [Ca^2+^]_SMC_ with the prescribed pressure *P*_MYOGENIC_ :

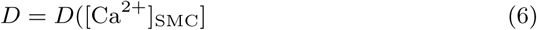

where the passive/active vessel mechanics, electrophysiology, and cross-bridge dynamics for the smooth muscles in the lumped arterioles are mostly from Carlson and Beard (2011) and Hai and Murphy (1998), and the others are enlisted in Table 2. In small resistant vessels, the perfusion follows the Poiseuille flow:

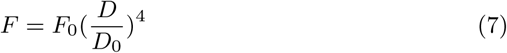

where the baseline diameter *D*_0_ and the corresponding perfusion flow rate *F*_0_ are prescribed for the condition without active force. The flow rate is proportional to the 4^th^ power of arterial diameter when the pressure difference is constant, i.e. very sensitive to the change of diameter.

**Table 2.** Smooth muscle and blood perfusion parameters.

| Symbol | Definition | Value |
| --- | --- | --- |
| $P_{\text{MYOGENIC}}$ | Arterioles transmural myogenic pressure | 100 mmHg |
| $\alpha_{\text{SMC}}$ | SMC plasma membrane area per unit volume | 0.32 / cm |
| $F_0$ | Perfusion constant | 0.7 mL/min/g |
| $D_0$ | Baseline diameter | 90 $\mu\text{m}$ |

**Table 3.**
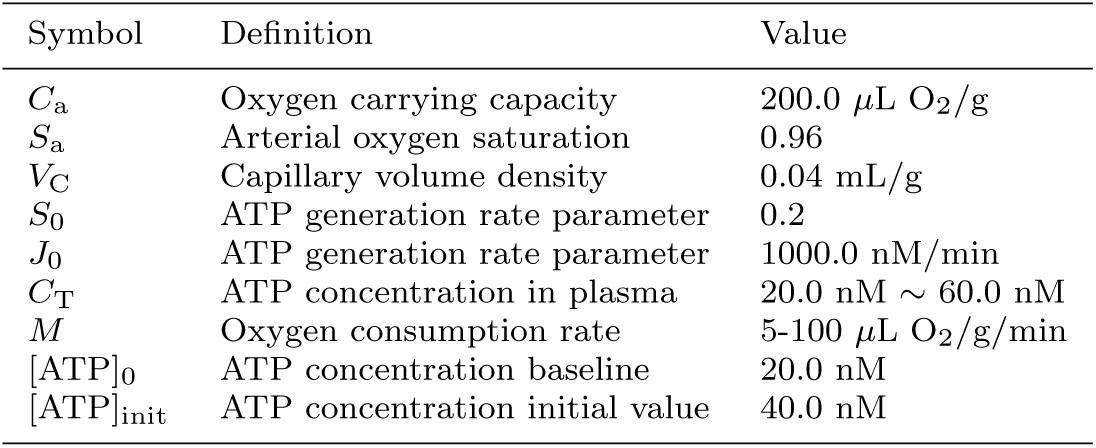
Oxygen sensing of ATP.

| Symbol | Definition | Value |
| --- | --- | --- |
| $C_a$ | Oxygen carrying capacity | 200.0 $\mu\text{L O}_2/\text{g}$ |
| $S_a$ | Arterial oxygen saturation | 0.96 |
| $V_C$ | Capillary volume density | 0.04 mL/g |
| $S_0$ | ATP generation rate parameter | 0.2 |
| $J_0$ | ATP generation rate parameter | 1000.0 nM/min |
| $C_T$ | ATP concentration in plasma | 20.0 nM ∼ 60.0 nM |
| $M$ | Oxygen consumption rate | 5-100 $\mu\text{L O}_2/\text{g}/\text{min}$ |
| $[\text{ATP}]_0$ | ATP concentration baseline | 20.0 nM |
| $[\text{ATP}]_{\text{init}}$ | ATP concentration initial value | 40.0 nM |

### 2.3 The oxygen sensing and ATP release model

The oxygen saturation in a capillary of length *L* is linearly dependent on the the steady-state oxygen consumption rate *M* :

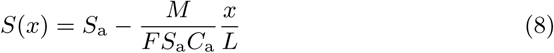

where *S*_a_ is the oxygen saturation in the arterioles, and *C*_a_ is the oxygen carrying capacity. Plasma ATP is governed by transport and production:

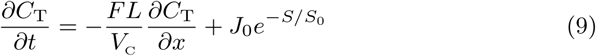

where *V*_C_ is the capillary volume density. Treated similar to Arciero et al. (2008), 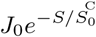 is the rate of ATP release from erythrocytes into plasma, exponentially decreasing depending on the oxygen saturation *S*.

### 2.4 Simulation protocol

The simulation protocol for an isolated EC tube (2 mm) is the following: (i) The direct/indirect activation of SK_Ca_ and IK_Ca_ channels via NS309/ACh continues from *t*=100 sec. (ii) From *t*=200 sec to 202 sec, an injection current 1nA is applied at Site 1. (iii) In case ACh is applied locally, the agonist is infused from *s*=100 to 500 *µ*m along the EC tube, and from *t*=100 sec to 150 sec in time. Also the external current 1nA is injected at *s*=250 *µ*m from *t*=133 sec to *t*=136 sec.

In the model capillary network simulation, we constructed five EC tubular segments (each 500 *µ*m) with two bifurcation nodes as shown in Figure 6(a). In the local zone of one bifurcation node, ACh in the amount of 6 *µ*M is infused for the whole simulation time. The current injection 5nA is applied for a finite time from *t*=20 sec to *t*=28 sec locally at an upstream site from a bifurcation node (Figure 6b).

**Fig. 1.**
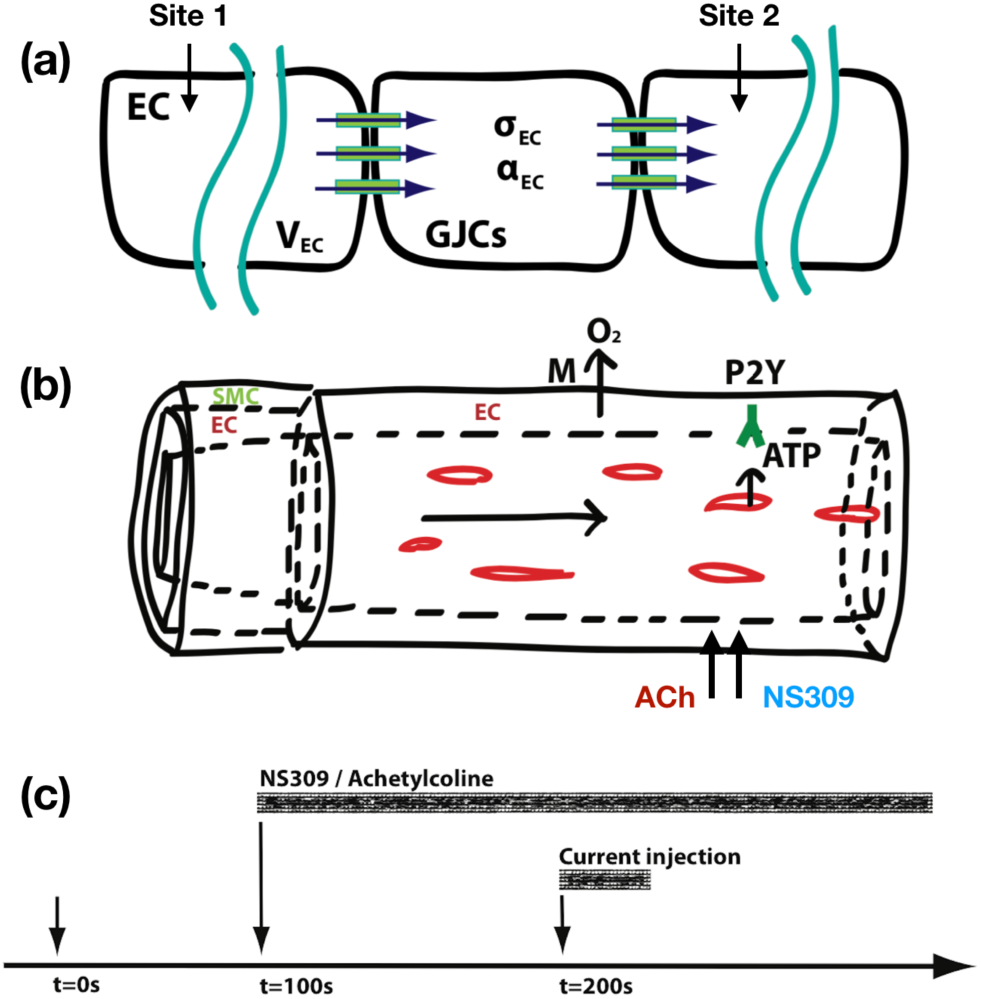
EC tube model for oxygen sensing and arterioles of endothelium coupled to smooth muscle. (a) The electrophysiology of endothelium is based on the model of Silva et al. (2009). The SK_Ca_/IK_Ca_ dependency on NS309 is newly included. The IP_3_ generation rate depends on the infused ACh or plasma ATP level. The cell-cell electrical interaction is through non-selective gap junctions. In the mono-domain formulation, *σ*_EC_ is the electrical conductivity in EC layer, and *α*_EC_ is the EC membrane area per unit volume. The electrical potential in the intracellular domain is defined as *V*_EC_. (b) The pre-capillary arterioles is basically composed of electrophysiology of endothelium (Silva et al. 2007) and myogenic smooth muscle (Carlson and Beard 2011). The oxygen consumption rate is denoted to *M*. (c) In the characterization of the EC tube model, NS309/ACh is infused from *t*=100 sec on and the current injections from *t*=200 sec to *t*=202 sec.

**Fig. 2.**
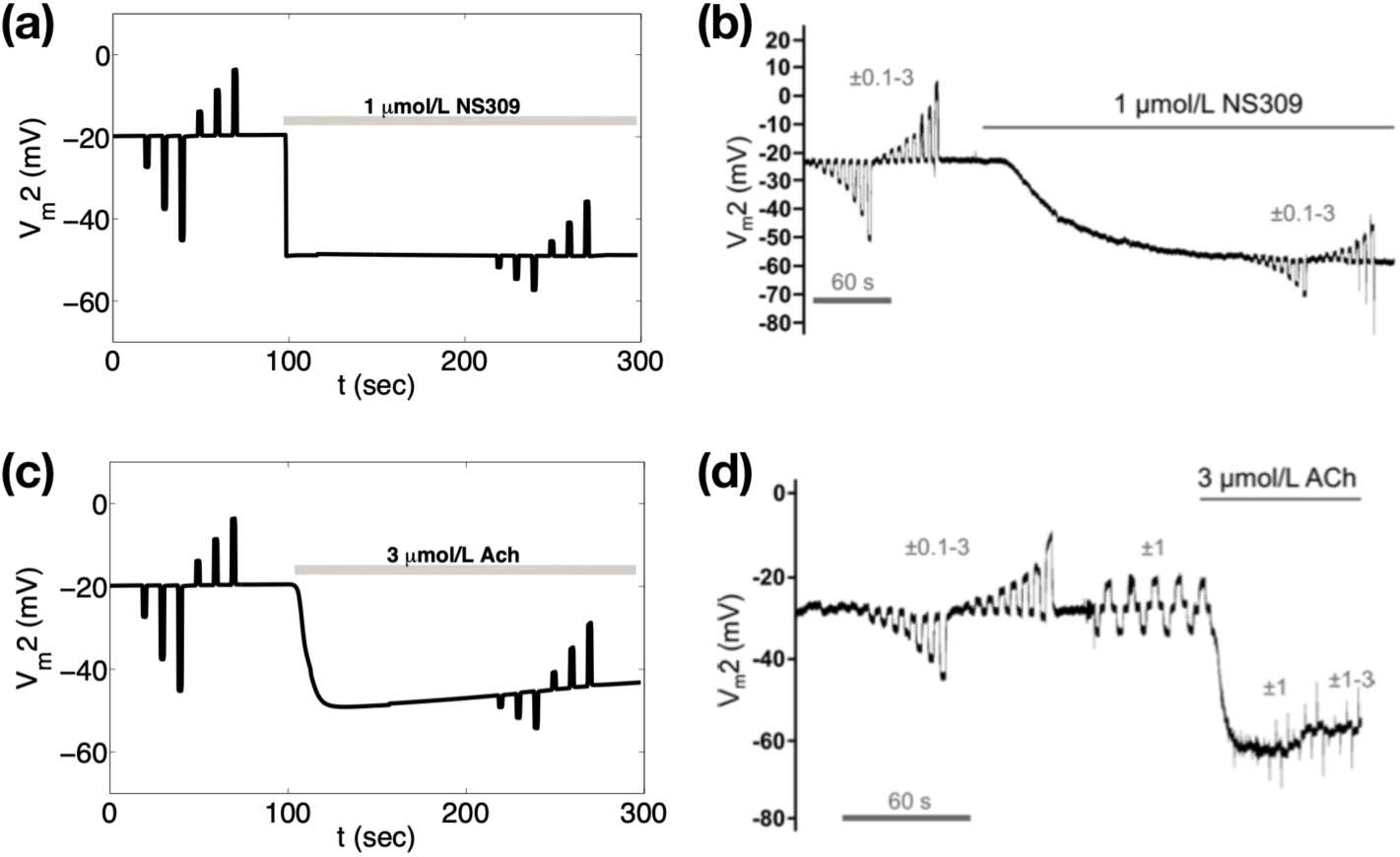
Time course of V_m_2 with NS309 and ACh. (a) Time course of *V*_m_ at *s* = 500 *µ*m (V_m_2) with 1 *µ*M NS309 from *t* = 100 sec on. Direct pharmaceutical activator NS309 activates directly SK_Ca_/IK_Ca_ and hyperpolarizes membrane potential to about −60 mV and attenuates response from current injection. EC tube model is simulated with current injection at Site 1 (1, 2, 3 nA) before and during NS309. (b) The corresponding experiment from Behringer and Segal (2012), Figure 2A. (c) Time course of *V*_m_ at *s*=500 *µ*m (V_m_2) with 3 *µ*M ACh from *t*=100 sec on. ACh activates indirectly SK_Ca_/IK_Ca_ via G-protein coupled receptors and hyperpolarizes membrane potential to about −60 mV and attenuates electrical conduction. Endothelial tube model is simulated with current injection at Site 1 (1, 2, 3nA) before and during ACh infusion. (d) The corresponding experiment from Behringer and Segal (2012), Figure 7A.

**Fig. 3.**
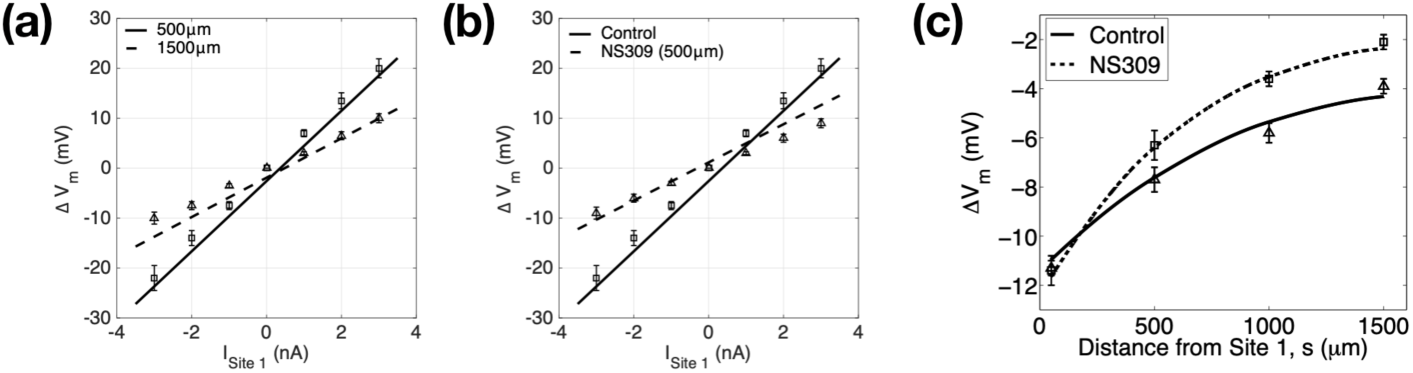
EC tube parameterization and validation. The mono-domain parameters (*σ*_EC_, *α*_EC_) are parameterized with the experimental data in Behringer and Segal (2012). (a) *Δ*V_m_2 at *s* = 500 *µ*m and *s* = 1500 *µ*m with current injection at Site 1 (1, 2, 3nA) (b) *Δ*V_m_2 at *s*=500 *µ*m with control (no NS309) and 1 *µ*M NS309. (c) *Δ*V_m_2 at *s*=50, 500, 1000, 1500 *µ*m with control (no NS309) and 1 *µ*M NS309.

**Fig. 4.**
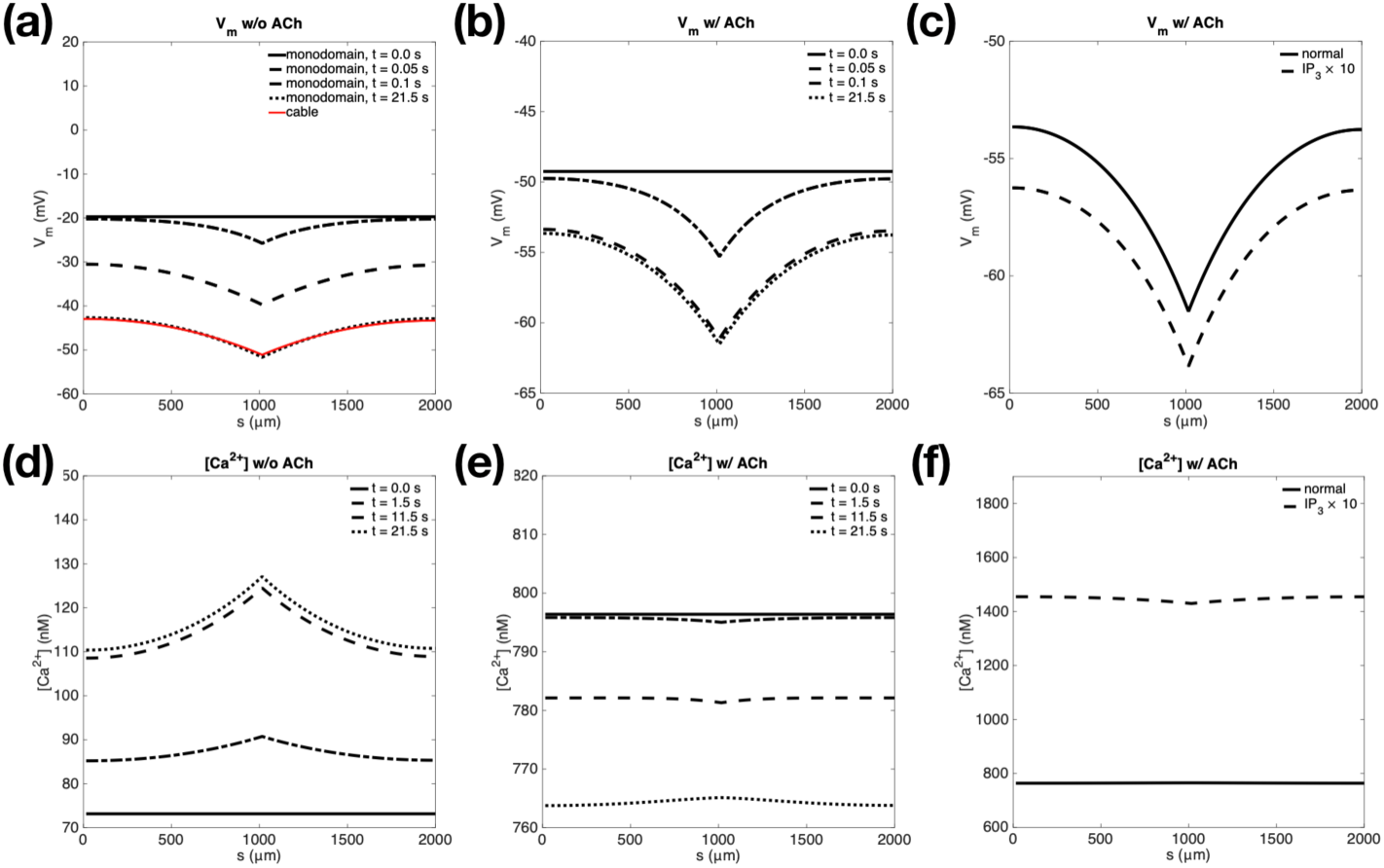
Mono-domain vs. cable equation and the effects of IP_3_R overexpression on Ca^2+^ dynamics. The current −3 nA is injected at *s*=1000 *µ*m from *t*=0 on. Endothelial layer is most hyperpolarized in the position of current injection, and the hyperpolarization is exponentially spread out. Accordingly the intracellular calcium is elevated with slower response than *V*_m_. (a) Without ACh, from resting potential *V*_m_ about −20 mV, the steady current injection hyperpolarizes EC tube to around −50 mV within 0.2 sec. The mono-domain electrical propagation is compared with that of cable 1D model (red curve). The involving parameters (*λ, r*_a_) are from the experimental data of Behringer and Segal (2012). (b) Cytosolic calcium is elevated from baseline level of 70 nM to ∼130 nM in 20 sec. (c) With ACh superfusion, the EC tube is hyperpolarized to −50 mV, and the membrane is more hyperpolarized to about −62 mV within 2.0 sec from the steady current injection. (d) At *t* = 0.0 sec, the calcium is already elevated from ACh activated IP_3_ induced calcium release, and the calcium level is lowered with calcium uptake and IP_3_ degradation. (e) and (f) With IP_3_R overexpression 10 folds, the positive feedback of calcium release from ER is strengthened, therefore cytosolic calcium gets higher, and the EC tube is more hyperpolarized.

**Fig. 5.**
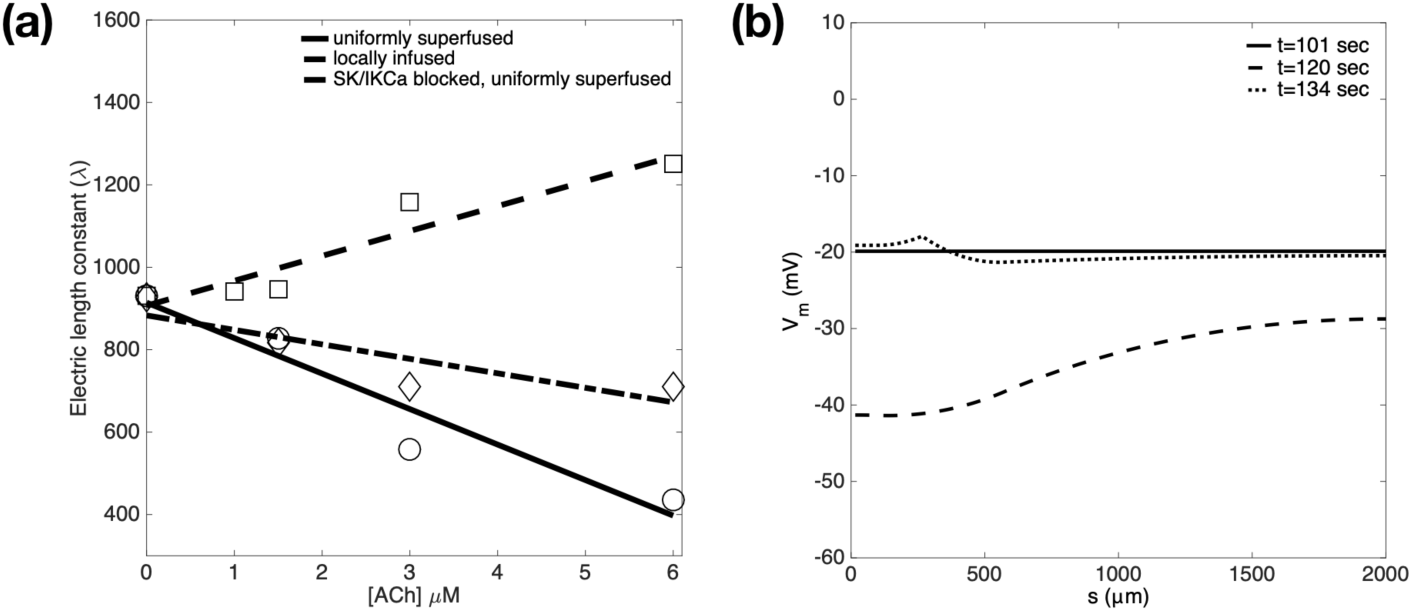
Electrical length constant vs. [ACh]. (a) The steady state IP3 generation rate 6.6 × 10^−5^ mM/s corresponds to [ACh] = 3 *µ*M with other values in a linear relationship. The electrical length constant is obtained from *ΔV*_m_2 at *s*=50, 500, 1000, 1500 *µ*m. In the solid curve with circle data points, the ACh is assumed to be uniformly superfused to the whole EC tube. The dotted curve with square data points represents the electrical propagation with local ACh infusion from *s* = 100 to 500 *µ*m along the tube, and from *t*=100 sec to *t*=150 sec. Also the current 1nA is injected at *s*=250 *µ*m from *t*=133 sec to *t*=136 sec. The dashdot curve is from SK_Ca_/IK_Ca_ channels blocked and EC tube uniformly superfused. (b) Spatial distribution at *t*=101, 120, 134 sec with local ACh infusion.

**Fig. 6.**
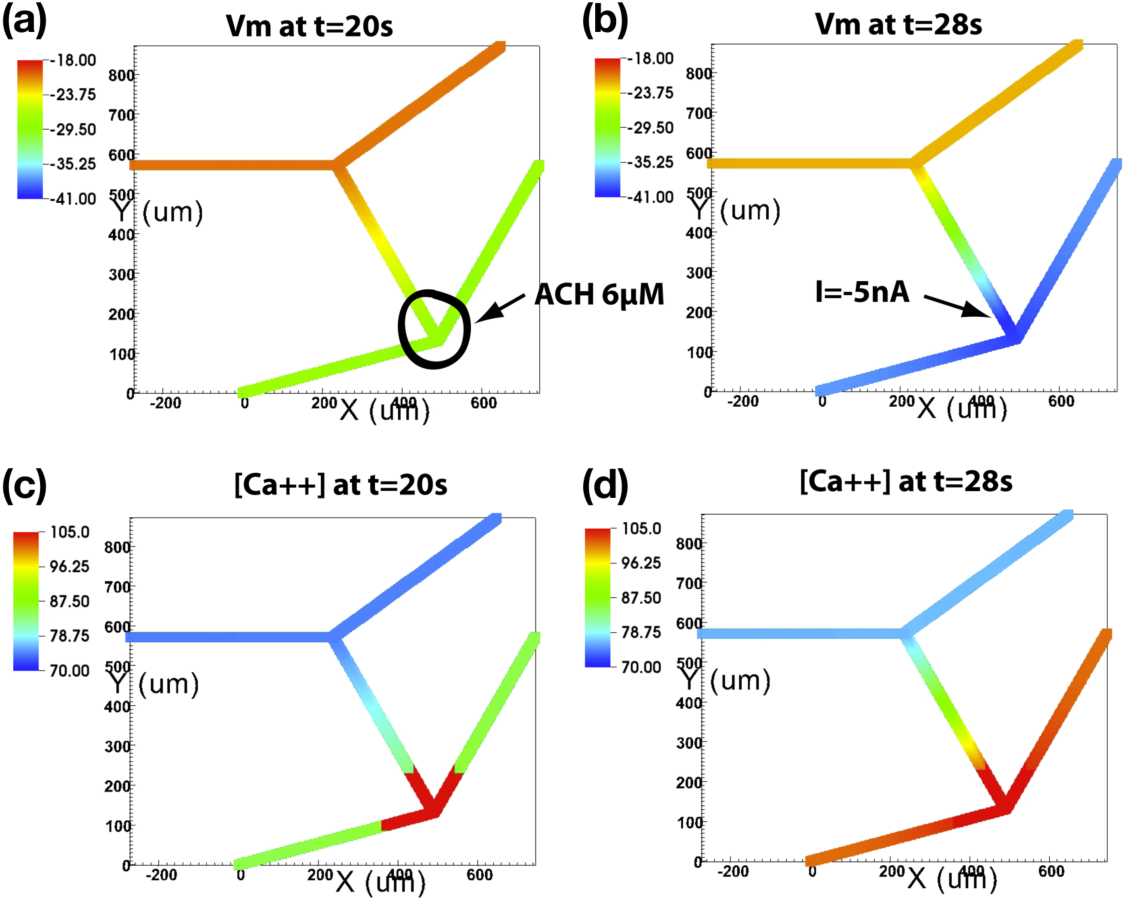
Propagation of hyperpolarization in a capillary model network. (a) In the whole simulation, 6 *µ*M ACh is locally infused in a local zone of one bifurcation node. (b) The current −5nA is injected at a position from *t*=20 sec to *t*=28 sec. Panels (a) and (b) show the spatial distribution of *V*_m_ at *t*=20 sec and *t*=28 sec; the propagated hyperpolarization from locally infused ACh, and further hyperpolarization from injected current to the upstream, respectively. (c) and (d) The spatial distributions of [Ca^2+^] at *t*=20 sec and *t*=28 sec; the local elevation of calcium level from the locally infused ACh, and the calcium wave propagation to the upstream with injected current, respectively.

For the oxygen sensing in the downstream, we focused on the isolated EC tube, and applied a range of oxygen consumption rate *M* (30-270 *µ*L O_2_/g/min). We measure the calcium level, membrane potential of smooth muscle cells in the lumped arterioles, and the dilated diameter and the perfusion flow rate.

To study the local infusion of ACh in the downstream of EC tube and see the vasodilation in the upstream, we have applied the ACh (6, 9, 12, 18, 24 *µ*M) from 1500 *µ*m to 2000 *µ*m, and measured the peak and relaxation of the dilatory response of the lumped arterioles.

### 2.5 Numerical methods

We solve the governing equations by the Krylov subspace iteration for the spatiotemporal propagation of the electrical potential using PETSc (Balay et al. 1997, Balay et al. 2019a,b). The computation on cellular solute/ionic transport is accelerated along the endothelial tube with parallelized GPU and Backward Differentiation Formula (BDF) using CVODE (Cohen and Hindmarsh 1996). For those solvers, absolute tolerance and relative tolerance are 10^−12^ and 10^−8^, respectively.

## 3 Results

As a first step, to parameterize the mono-domain model, we used the electrical propagation of endothelial tube from experimental data of skeletal muscle (Behringer and Segal 2012). As described in the Simulation Protocol (Figure 1c), the activation/inactivation of SK_Ca_ and IK_Ca_ continues from *t*=100 sec on. The direct activation of SK_Ca_/IK_Ca_ are realized by treating 1 *µ*M NS309, and the indirect activation by 3 *µ*M ACh. The current is injected at Site 1 from *t*=200 sec to *t*=202 sec. Figure 2(a) and (c) show the simulated time courses of *V*_m_ at the distance *s*=500 *µ*m from Site 1 when NS309 and ACh are superfused uniformly through the whole EC tube domain, respectively. When SK_Ca_ and IK_Ca_ are activated, the endothelial tube is hyperpolarized from the resting state about −20 mV to −60 mV. In control with SK_Ca_ and IK_Ca_ inactivated before *t*=100 sec, the endothelial tube stays in the initial resting potential, and the maximum peaks of *ΔV*_m_ are larger in comparison to the responses in hyperpolarized states. Figure 2(b) and (d) correspond to the experimental data from the application of the pharmaceutical drug NS309 and the endogenous agonist ACh (Behringer and Segal 2012).

Figure 3 shows the comparison between the experimental data of electrical signal propagation on the EC tube and the model simulation with two primary parameters, the endothelial electrical conductivity *σ*_EC_ and the endothelial plasma membrane area per unit volume *α*_EC_. In the parameterization, we have used the electrical propagation data with activated/inactivated SK_Ca_ and IK_Ca_. The data used in the parameterization is *ΔV*_m_, the maximum difference of membrane potential before and during membrane depolarization after current injection. Those *ΔV*_m_ are collected from *s*=50, 500, 1000, and 1500 *µ*m. Figure 3(a) shows the linear curve fitting with graded current injection (1, 2, 3 nA) at *s*=500 and 1500 *µ*m in control. The solid curve is from simulated *ΔV*_m_ at *s*=500 *µ*m and the corresponding square data points/error bars are from the experiments. The dotted curve is from simulated *ΔV*_m_ at *s*=1500 *µ*m and the corresponding triangle data points/error bars are from the experiments. With the electrical propagation attenuation, the slope of the curves is decreased. Figure 3(b) demonstrates the linear curve fitting with *ΔV*_m_ at the same position *s*=500 *µ*m with NS309 (1 *µ*M) and control (before NS309 treatment, *s*=500 *µ*m). The solid curve is from simulated *ΔV*_m_ at control and the corresponding square data points/error bars are from the experiments. The dotted curve is from simulated *ΔV*_m_ with NS309 treatment and the corresponding square data points/error bars are from the experiments. With the hyperpolarization of the EC tube, the slope of the curves is decreased. In Figure 3(c), we fitted the electrical signal attenuation with different current injections (1 nA for control, 2 nA for NS309) from a couple of distances, *s*=50, 500, 1000, 1500 *µ*m. The solid curve is from simulated *ΔV*_m_ at control and the corresponding triangle data points/error bars are from the experiments. The dotted curve is from simulated *ΔV*_m_ with NS309 treatment and the corresponding square data points/error bars are from the experiments. The parameterized EC tube model satisfies quantitatively the electrical length constants in the control and NS309 application.

From the parameterized EC tube model, we investigated the effects of ACh in the electrical propagation and calcium dynamics. In Figure 4, we show the spatial distribution of *V*_m_ and [Ca^2+^] with and without ACh. When the agonist is not applied, the resting potential stays at around −20 mV, and the intracellular calcium level is around the baseline of 70 *µ*M. However, when ACh is infused, the membrane is hyperpolarized, and the intracellular calcium level arises to about 795 *µ*M, and the membrane potential goes down to about −50 mV. In a way of model validation, this is compared with the profile from 1D cable equation (Figure 4a, red curve). The involving parameters (*λ, r*_a_), the electrical length constant and axial resistance to current flow are from the experimental data of Behringer and Segal (2012). In the spatial distribution, *V*_m_ is at the peak in the position of current injection, and exponentially spreads out. The response of *V*_m_ is fast so that after the current injection is finished at *t*=2.0 sec, *V*_m_ gets back to the initial resting potential about −20 mV within 2.5 sec. In contrast, the calcium response is slow with no change in calcium level at *t*=0.25 sec after current injection at *t*=0.0 sec. At *t*=2.5 sec, the intracellular calcium is still recovering in transition whether the EC tube is infused with ACh or not.

Figure 5 shows the electrical length constant *λ* versus infused ACh concentration. This parameter was obtained from *ΔV*_m_2 at *s*=50, 500, 1000, and 1500 *µ*m similar to the data in Figure 3(c). We observed opposite trends depending on the spatial range of infused ACh. When the ACh is locally infused, the electrical length constant increased 35% with ACh infusion up to 6 *µ*M, but *λ* gets short to about 50% when ACh is globally perfused. Blocking SK_Ca_ and IK_Ca_ channels suppresses the attenuated electrical propagation (turning less stiff from solid to dashdot profile in Figure 5a).

Next, to realize the electrical signal propagation in microvascular network, we observed the ascending hyperpolarization on a model network with each segment 500 *µ*m (Figure 6). In the whole simulation, an amount of ACh (6 *µ*M) is locally infused in the region of one bifurcation node as indicated in Panel (a). Also a local current −5 nA is injected at a position indicated in Panel (b) from *t*=20 sec to *t*=28 sec. Panels (a) and (b) show the spatial distribution of *V*_m_ at *t*=20 sec and *t*=28 sec. Panel A shows the spreading hyperpolarization from locally infused ACh, and Panel (b) demonstrates further hyperpolarization to upstream branches from injected current. Panels (c) and (d) show the spatial distribution of [Ca^2+^] at *t*=20 sec and *t*=28 sec. Panel (c) shows the local elevation of calcium level from the locally infused ACh, and Panel (d) illustrates the calcium wave propagated to the upstream with the injected current.

Now, we investigate the influence of local infusion of ACh in the terminal area of EC tube, and how the post-capillary arterioles respond in vasodilation. ACh in the amount of 6, 9, 12, 18, 24 *µ*M are applied in the downstream from 1500 *µ*m to 2000 *µ*m. The peak of the dilatory response of the lumped arterioles is shown in Figure 7(a). Interestingly there is a biphasic profile in maximum response from local ACh infusion, with the maximum vasodilation about 17% with 12 *µ*M of ACh, which is comparable to the experiments in Doyle and Duling (1997). The relaxation time from dilation to the resting state gets short from about 500 sec for ACh 6 *µ*M to 30 sec for ACh 24 *µ*M.

**Fig. 7.**
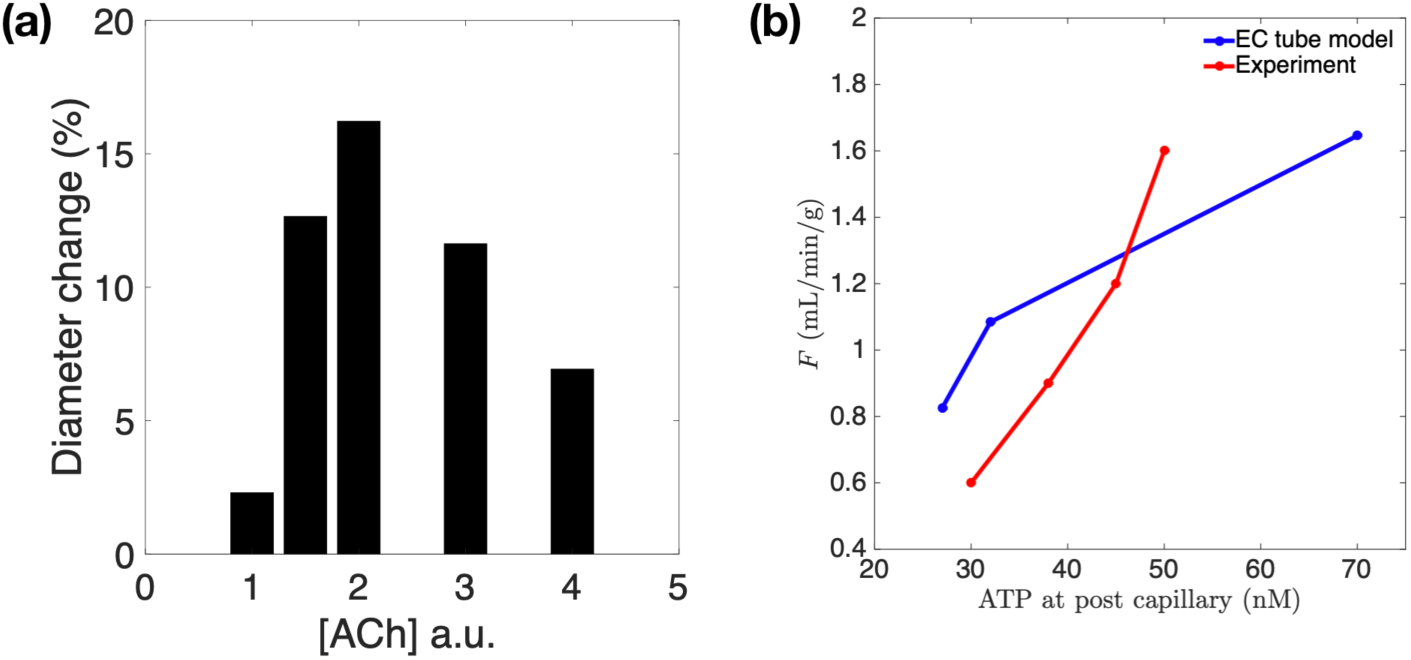
(a) The maximum vasodilatory increment versus local infusion of ACh (b) Blood flow density versus plasma ATP in terms of oxygen consumption rate. ATP at the post capillary domain versus pre capillary blood flow density. The EC tube model of ascending vasodilation is compared with coronary blood flow regulation of Farias et al. 2004

Finally, we characterize the influence of oxygen consumption and induced oxygen tension for ATP activation and the ascending vasodilation. We have prescribed the baseline oxygen consumption *M*_0_ with 30 *µ*L O_2_/g/min, and considered two cases of *M* with 80 and 270 *µ*L O_2_/g/min. Figure 8(a) shows the membrane potential distribution. With the oxygen consumption increment, the plasma ATP level is increased (Figure 8e), and through purinergic signaling, intracellular calcium level is also elevated from purinergic IP_3_ generation increment and calcium release from ER (Figure 8c). Finally from calcium-activated potassium channels, the membrane potential is more hyperpolarized (Figure 8a). Figure 8(b), (d), and (f) demonstrate the oscillation of membrane potential, calcium concentration at the terminal downstream with the increased oxygen consumption.

**Fig. 8.**
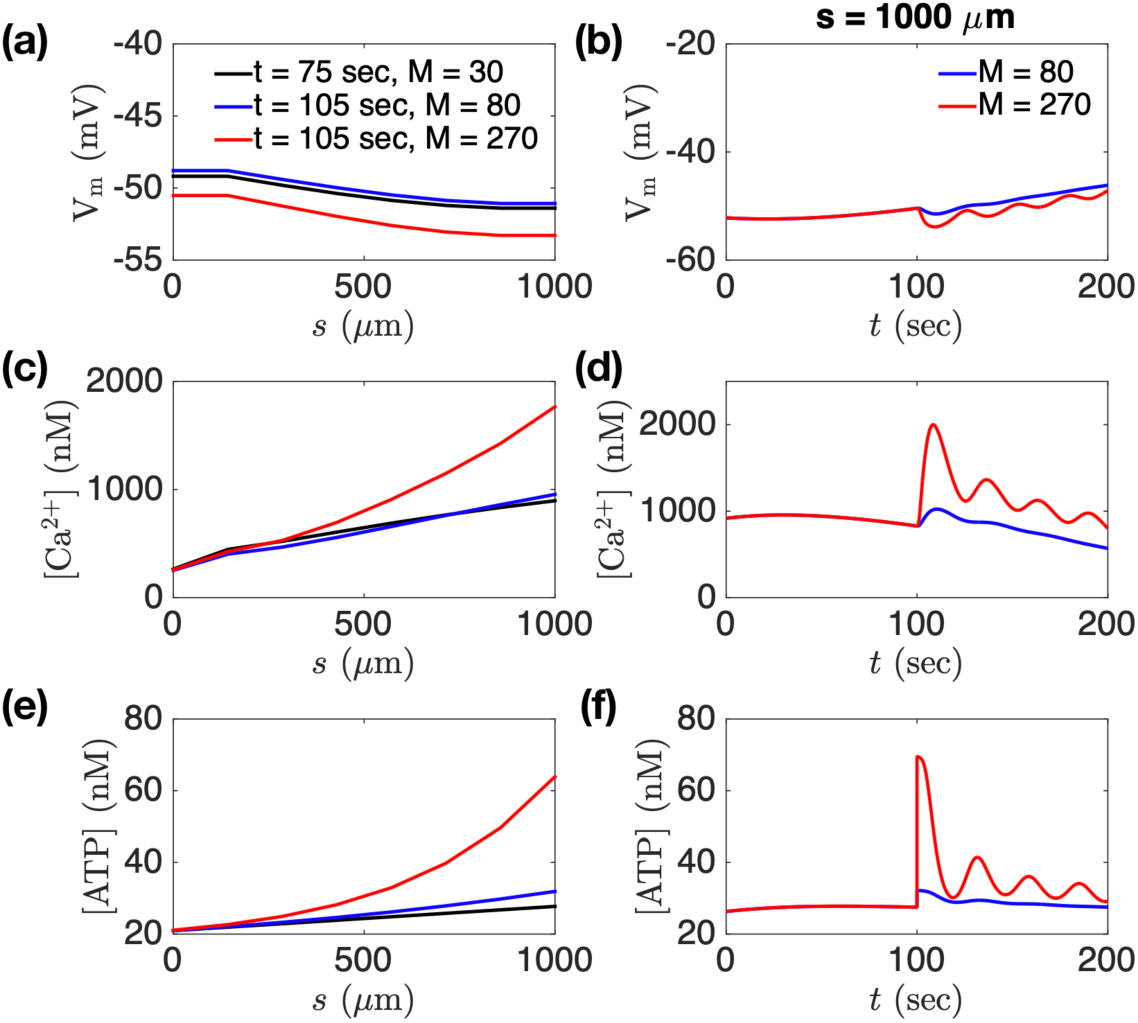
Endothelial oxygen sensing dependent on oxygen consumption rate *M*. (a) and (b) Membrane potential distribution. With the oxygen consumption elevation, the membrane is hyperpolarized. (c) and (d) Calcium concentration distribution along the tube. With the oxygen consumption elevation, the intracellular calcium level is elevated from purinergic IP_3_ generation increment and calcium release from ER. (e) and (f) Plasma ATP level is increased with the oxygen consumption elevation, the low oxygen tension in post-capillary domain, and the ATP release from red blood cells. The oscillatory profiles are emerging when the oxygen consumption is highly elevated to 270 *µ*L O_2_/g/min.

Figure 9 illustrates the time courses in electrophysiology from the SMC in the arterioles and vessel dilation. When the downstream oxygen consumption is surged up from *M* =80 to *M* =270 *µ*L O_2_/g/min, the hyperpolarized endothelial membrane potential is propagated to the upstream arterioles, and decreases the intracellular calcium concentration of SMC (Figure 9a). The smooth muscle in the upstream arterioles shows oscillatory profiles in IP_3_ and calcium concentrtions, and the membrane potential. Here, the oscillatory period is around 25 sec. During 100 sec after *M* is increased to 270 *µ*L O_2_/g/min, via this vasomotion, the capillary domain is thought to be more oxygenated and ATP level is gradually decreased recovering the resting state. Figure 7(b) shows ATP at the post capillary domain versus pre capillary blood flow density in terms of oxygen consumption rate, and this is compared with the coronary blood flow regulation of Farias et al. (2005). In the experiment, during exercise, the myocardial oxygen consumption increased about 3.2 times, and coronary blood flow increased about 2.7 times. Accordingly the oxygen tension decreased from 19 to 12.9 mmHg, and coronary venous plasma ATP increased from 31.1 to 51.2 nM. The current model captures this trend qualitatively.

**Fig. 9.**
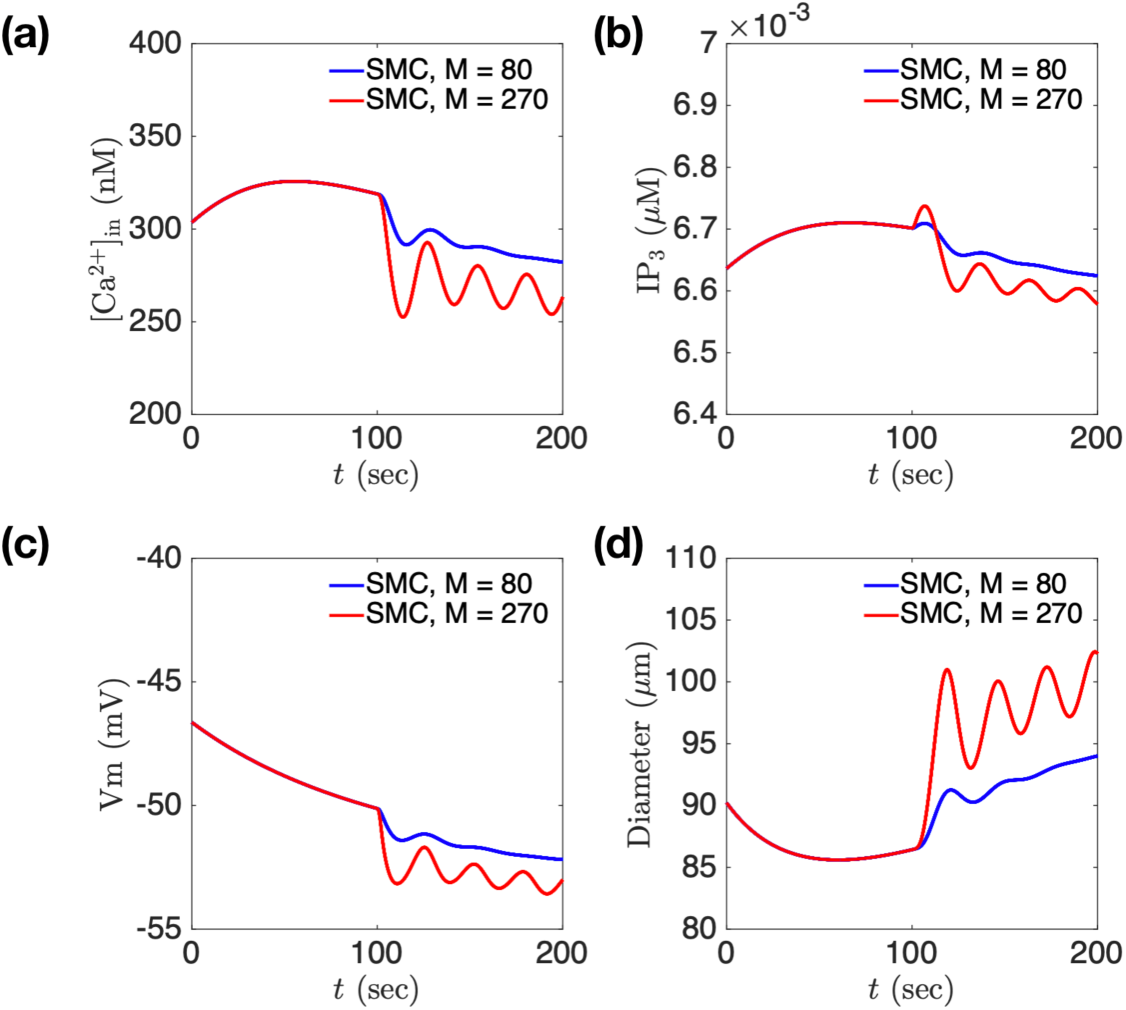
Arterioles SMC time course in electrophysiology and vessel dilation. Until *t* = 100 sec, *M* is set to 30 *µ*L O_2_/g/min. From *t*=100 sec, *M* is set to 80/270 *µ*L O_2_/g/min. Enhanced oxygen consumption induces vasomotion. (a) Intracellular calcium concentration is decreased while transiting from non-oscillating to oscillating state from oxygen consumption rate increment. (b) IP_3_ level is decreased while non-oscillating/oscillating from oxygen consumption rate increment. (c) Membrane is further hyperpolarized while non-oscillating/oscillating from oxygen consumption rate increment. (d) Smooth muscle is dilated without and with vasomotion when a tissue extracted oxygen further for the blood flow increase.

## 4 Discussion

The proposed mono-domain model captures quantitatively the electrical propagation along the EC tube in control, and also satisfies the attenuated electrical propagation from hyperpolarization. The proposed integrated model of EC tube and lumped arterioles vessel mechanics shows the upstream response of vasodilation from the localized agonists of ACh and ATP.

The membrane depolarizes and repolarizes quite quickly in response to the current injection within a couple of seconds (Figure 2), that is realized by changing the EC tube membrane capacitance as a low value in comparison to the cellular model in Silva et al. (2009). Here one thing that stands out is the response to NS309 stimulation. In our model, NS309 application causes an immediate hyperpolarization. However, the data show a much slower trend. We speculate that in the experiment, the drug is slow to get to the site of action, possibly on the inside of the EC. The process of calmodulin domain sensitization and slow calcium-activated potassium channel activation to resting levels of intracellular calcium may account for the delay. With NS309 (and no significant increase from resting calcium), SK_Ca_/IK_Ca_ activation relies upon resting calcium levels despite a large shift in calcium sensitivity. Presumably, this would take a very long time relative to activation via ACh (massive increase from resting calcium) where calcium ions are readily available for SK_Ca_/IK_Ca_ activation. However, this delay of hyperpolarization to NS309 appears to decrease with higher doses.

ACh works as vasodilator in the microvasculature, and some experiments show that the application of ACh enhances the ascending hyperpolarization and increases electrical length constant (Emerson et al. 2002). In one protocol of the current model simulations, we show an opposite trend that ACh attenuated the electrical propagation (Figure 5). One reason for this discrepancy is thought to be from different experiment/simulation setup. In the experiment, the perfusion of ACh is applied in a local region. On the other hand, the one specific model simulation implements the ACh uniformly in the whole EC tube. While interrogating this opposite trend on the ACh dependency, we also simulated with local application of ACh, and indeed, increased up to 35% with the agonist level from 0 to 6 *µ*M. In the uniformly perfused ACh in the whole EC tube, more ACh evokes more hyperpolarization, and the depolarization from current injection is suppressed. This is the primary mechanism for further attenuation from higher ACh. In contrast, when ACh is infused locally, more ACh induces deeper hyperpolarization, and the membrane voltage gradient between ACh infused zone and ACh free zone gets bigger, and the electrical currents are increased through the ACh transitional area. This is supposed to drive increased electrical conduction and prolonged *λ* from higher acetylcholine.

The simulation of model capillary network shows the electrical and calcium signal propagation in the scale of ∼1 mm, and indeed in the branched network. We have shown the propagation of hyperpolarization from the local perfusion of ACh (Figure 6a), and highlighted ascending electrical propagation to the upstream from a local current injection (Figure 6b). In our modeling, we did not include diffusion terms for calcium and IP_3_, i.e. the transport of calcium and IP_3_ through gap junctional channels, which may enhance the the electrical propagation somehow, but this is very slow in comparison to the electrical effects (Figure 4). The calcium wave propagation in Figure 6(c) and (d) is purely from electrical hyperpolarization.

The current model does support the ATP hypothesis (Gorman et al. 2010) by the proposed mechanistic integrated model. Interestingly, we recapitulate vasomotion in the lumped arterioles and the corresponding oscillatory profiles in ATP and calcium dynamics in the downstream EC tube. Vasomotion is thought to enhance the perfusion and oxygenation by blood circulation to the capillary domain (Goldman and Popel 2001, Domeier and Segal 2007, Thorn et al. 2011, Haddock et al. 2011). In the current induction of oscillation in the arterioles and EC tube, the oscillatory frequency is around 0.04 Hz. In the brain, low-frequency calcium oscillation related to cerebral hemodynamics is about 0.07 Hz in rat somatosensory cortex (Du et al. 2014). In the hemodynamics of skeletal muscle with local femoral pressure reduction, there comes a slow-wave flow motion with 0.025 Hz (Schmidt et al. 1992). The modeling study of renal myogenic autoregulation and vasomotor response reconstructs 0.1-0.2 Hz vasomotion based on rat afferent arteriole (Sgouralis and Layton 2012). We expect to do more rigorous analysis for characterizing precise conditions generating vasomotion from multiple factors (Sgouralis and Layton 2012, Kapela et al. 2012, Arciero and Secomb 2012) including internal states of pressure and oxygen consumption as well as the amounts of endogenous and pharmaceutical agonists. All together, this is indeed an integrated bidirectional feedback between the vasodilatory perfusion in the upstream and the oxygen sensing associated ATP release in the downstream, mediated by the ascending electrical signal along capillary networks.

For the future work, we remain the electrical propagation in realistic capillary networks, and oxygen consumption in specific tissues with specific conditions not only of physiological states, but also pathphysiological states of ischemia, hypertension, and metabolic syndromes. We can easily apply localized perfusion of acetylcholine and pharmaceutical vasodilators in the presence of diverse forms of endothelial dysfuncition in the network in order to investigate the global therapeutic effectiveness of vasodilation and oxygenation.

## Acknowledgements

The author acknowledges the valuable discussion in modeling with Ranjan Pradhan, Brian E. Carlson, and Daniel A. Beard. Especially the formulation of the open probability in SK_Ca_ and IK_Ca_ with NS309 is accredited to Ranjan Pradhan. The author also thanks to Steven Segal and Erik Behringer for the experimental data and helpful discussion. This research is partially supported by U01 HL118738-01A1 and NIGMS-P50GM094503.

